# Automated literature mining and hypothesis generation through a network of Medical Subject Headings

**DOI:** 10.1101/403667

**Authors:** Stephen Joseph Wilson, Angela Dawn Wilkins, Matthew V. Holt, Byung Kwon Choi, Daniel Konecki, Chih-Hsu Lin, Amanda Koire, Yue Chen, Seon-Young Kim, Yi Wang, Brigitta Dewi Wastuwidyaningtyas, Jun Qin, Lawrence Allen Donehower, Olivier Lichtarge

## Abstract

The scientific literature is vast, growing, and increasingly specialized, making it difficult to connect disparate observations across subfields. To address this problem, we sought to develop automated hypothesis generation by networking at scale the MeSH terms curated by the National Library of Medicine. The result is a Mesh Term Objective Reasoning (MeTeOR) approach that tallies associations among genes, drugs and diseases from PubMed and predicts new ones.

Comparisons to reference databases and algorithms show MeTeOR tends to be more reliable. We also show that many predictions based on the literature prior to 2014 were published subsequently. In a practical application, we validated experimentally a surprising new association found by MeTeOR between novel Epidermal Growth Factor Receptor (EGFR) associations and CDK2. We conclude that MeTeOR generates useful hypotheses from the literature (http://meteor.lichtargelab.org/).

**AUTHOR SUMMARY:** The large size and exponential expansion of the scientific literature forms a bottleneck to accessing and understanding published findings. Manual curation and Natural Language Processing (NLP) aim to address this bottleneck by summarizing and disseminating the knowledge within articles as key relationships (e.g. TP53 relates to Cancer). However, these methods compromise on either coverage or accuracy, respectively. To mitigate this compromise, we proposed using manually-assigned keywords (MeSH terms) to extract relationships from the publications and demonstrated a comparable coverage but higher accuracy than current NLP methods. Furthermore, we combined the extracted knowledge with semi-supervised machine learning to create hypotheses to guide future work and discovered a direct interaction between two important cancer genes.

## INTRODUCTION

It is difficult to keep abreast of new publications. Currently, PubMed contains over 28 million papers (http://www.ncbi.nlm.nih.gov/pubmed)—3 million more than three years ago. This steady accumulation of findings gives rise to a large number of latent connections that Literature-Based Discovery (LBD) seeks to systematically recognize and integrate [1], such as Swanson’s original finding linking fish oil to the treatment of Raynaud’s disease [2]. Since this original analysis, LBD has been extensively replicated, automated and expanded [3-10], leading to new patterns of inference – e.g. locating opposing actions of a disease and a drug on given physiological functions [11] – and to new discoveries [12]. Successes include the automated discovery of protein functions [13, 14] and of the genetic bases of disease [15, 16], as well as the stratification of patient phenotypes [17] and outcomes [18].

A limitation of LBD, however, is its dependence on knowledge extraction. It either relies on human curation, which is not scalable, or on comprehensive text-mining, for which algorithms are less accurate [19, 20]. One of the largest curated multi-modal biomedical data sources is the Comparative Toxicogenomics Database (CTD). CTD relied on five full-time biocurators to curate 70-150 articles a day [21] and gather drug-gene, drug-disease, and gene-disease associations from 88,000 articles, or about 0.3% of PubMed. By contrast, Natural Language Processing (NLP) combines semantic analysis of word meaning with syntactic knowledge of word grammar to break down sentences into biomedical associations. It automatically extracts knowledge from the entire literature without human supervision [22, 23], and it is improving [24] but still much less accurate than human curation [23, 25].

To combine the benefits of human curation with the scalability of text-mining, we note that an exhaustive manual curation of PubMed articles already exists. In order to facilitate article indexing and retrieval, curators at the National Library of Medicine assign Medical Subject Headings (or MeSH terms) and Supplemental Concept Records (SCR) to every PubMed article. These terms (https://www.nlm.nih.gov/pubs/factsheets/mesh.html) summarize key biomedical concepts for each paper, and to expand coverage and refine relevance, they are revised annually (or daily for SCRs) [26] (https://www.nlm.nih.gov/pubs/factsheets/mesh.html). The co-occurrence of MeSH terms with text-mined gene names was used to cross-reference genes and predict diseases that shared disease characteristics and chromosomal locations [27, 28].

Unfortunately, this was dependent on NLP for the identification of the genes (due to a reported low-coverage of gene MeSH terms in 2003) and required additional databases of information for chromosomal locations. Another study suggested that weighting MeSH terms (TF*IDF) was beneficial [29]. More recently, MeSH term co-occurrence was analyzed with various unsupervised and supervised techniques to make retrospective and prospective hypothesis [30] that predicted future associations between MeSH terms accurately [30]. This approach used all MeSH terms, including broad terms such as “Proteins”, but not SCR. Unfortunately, the individual terms were not mapped to canonical gene and drug terms, such as HGNC[31] and PubChem [32] identifiers restricting comparisons to curated datasets. Overall, the use MeSH terms in LBD has been limited in a few applications with regards to gene accuracy/coverage, selection and mapping of MeSH terms, and comparisons to curated datasets.

To improve on the generality, scalability and accuracy of these approaches we sought to comprehensively use MeSH terms for genes, to add the information from SCRs, and to perform thorough comparisons against biological standards and among the latest NLP methods. We also developed a robust unsupervised link prediction algorithm and experimentally tested a top prediction. The result is a literature-derived network called MeTeOR (the MeSH Term Objective Reasoning approach), which represents gene-drug-disease relationships exclusively from MeSH term and SCR co-occurrence. We show below that MeTeOR supplements knowledge from reference databases and more accurately recovers known relationships than traditional text-mining. Pairing the MeTeOR network with Non-Negative Matrix Factorization (NMF), an unsupervised machine learning algorithm, significantly improved LBD performance.

## RESULTS

### Developing a literature-based network from MeSH terms

In order to represent published biological associations among genes, drugs, and diseases, we took the Medical Subject Headings (MeSH) and Supplemental Concept Records (SCR) assigned to more than 21,531,000 MEDLINE articles by the National Library of Medicine (NLM) (**Supplemental Fig. 1**). MeSH terms facilitate indexing and searching, and SCR terms were created to identify drugs too numerous to be directly added as MeSH terms. (SCR terms also represent diseases and genes, among other topics.) Each distinct MeSH and SCR term became one of 276,000 nodes with 286 million term-article relationships. Nodes that co-occurred in a paper were fully connected into a clique for each article, and cliques were joined when they shared nodes across articles (**Figure 1A**). This generated a single network with 129 million term-term non-overlapping edges in which the number of articles that gave rise to a given pair of nodes measures the confidence of their association. Of these nodes, 39% mapped to 89,000 drugs, 4,800 diseases, and 13,000 genes, forming 9 million edges. The network consisted primarily of genes (12%) and drugs (82%), but, given the focus of much biomedical research on disease, 56% of edges contained a disease (**Figure 1B**). As articles get added to MEDLINE, the network can be updated as soon as they have been annotated by the NLM.

**Figure 1.**
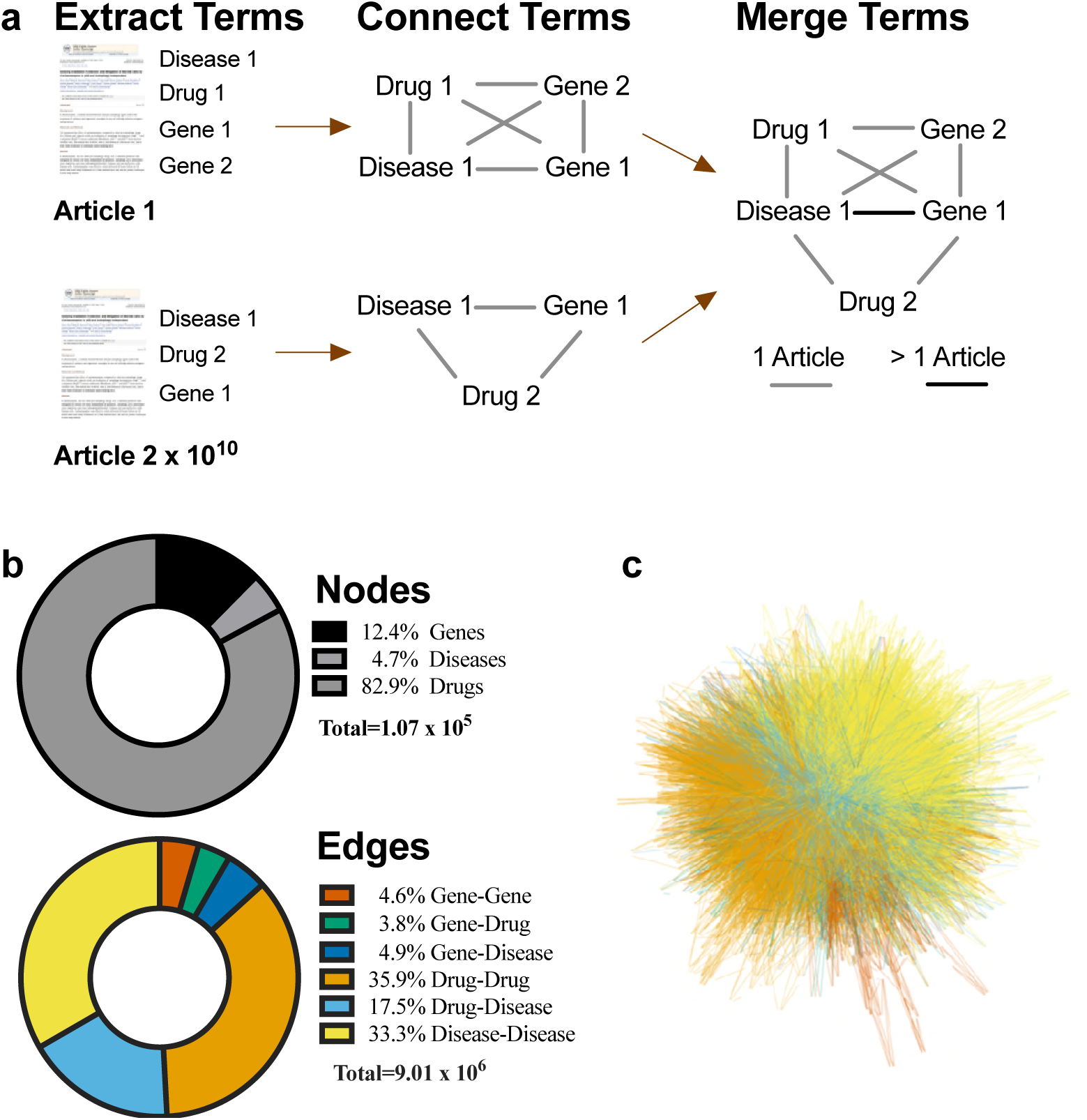
MeSH terms can provide a reliable approximation of biomedical knowledge in the literature. A) MeSH terms are taken from an article, connected into a clique, and then merged by nodes across over 22 million articles in PubMed. Any associations that overlap between articles are considered to have greater confidence. B) Graphical representation of the proportion of nodes for each entity type and the percentage of edges per association type. C) The MeTeOR network as a whole is formidable, despite the exclusion of all edges with a confidence of less than 200 articles.

This network was too visually dense to interpret, even when focusing on only high-confidence relationships (conf. >200 articles, degree >3) (**Figure 1C**). The complexity of the network and the presence of complete cliques at the article-level led us to evaluate the network’s topology. When limited to genes, drugs, and diseases, MeTeOR best fits a scale-free network with a power-law distribution of node degrees, where γ ≈ 1.34 (*p*-value ≪ 10^-35^ compared to log-normal and exponential distributions; **Supplemental Fig. 2**) [33] and some nodes have a much higher node degree, i.e. greater connectivity. The presence of such hubs is a common feature of real-world networks [34]. MeTeOR thus condenses PubMed knowledge into a computable and well-structured network that is amenable to analysis by established network algorithms.

**Figure 2.**
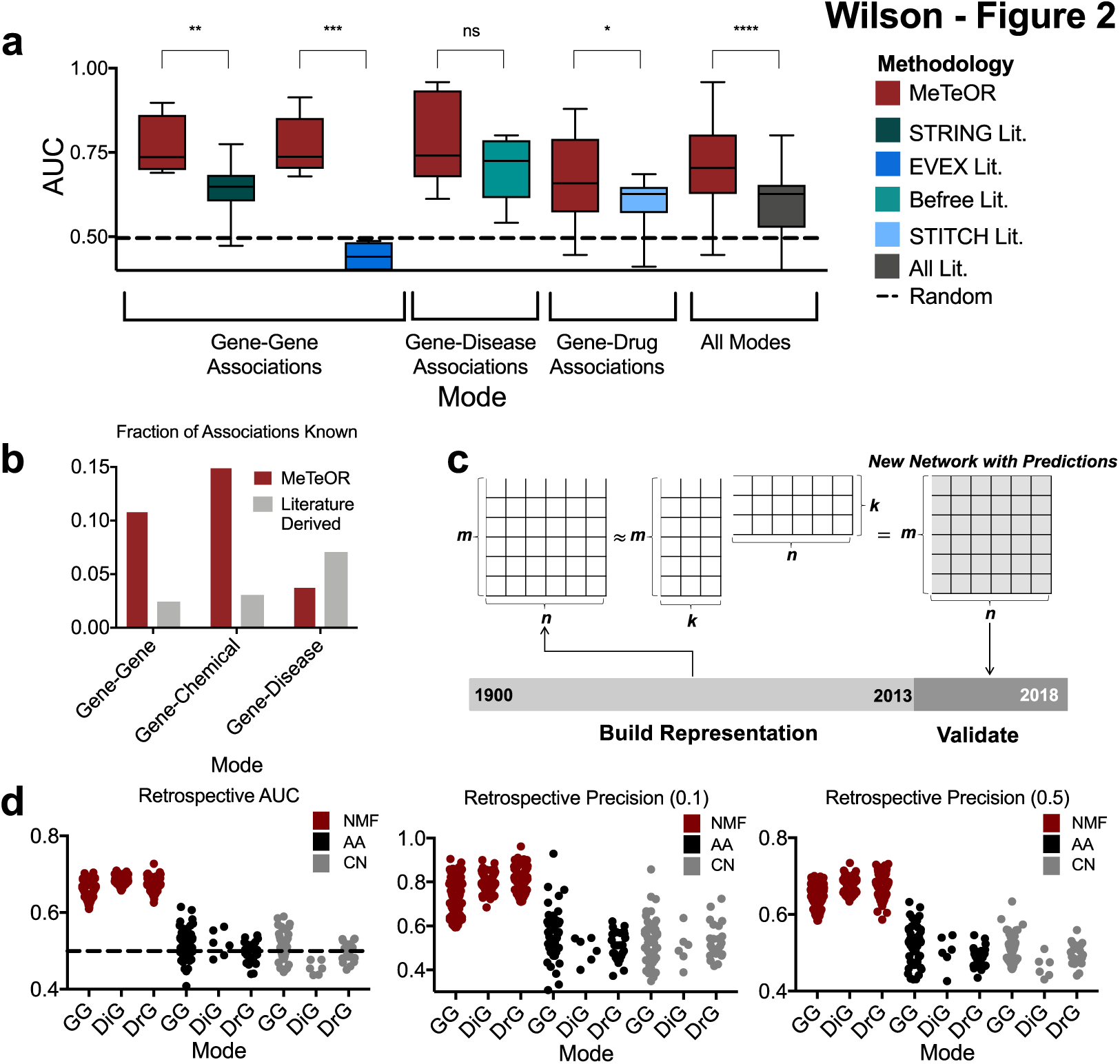
MeTeOR reliably recovers known associations and predicts new biomedical knowledge. A) MeTeOR was compared with appropriate algorithms using the area under the Receiver Operating Characteristic (ROC). All tests were done with bootstrapping to give a confidence range and to balance the positives and negatives. The literature-derived algorithms were STRING-Literature, EVEX, STITCH-Literature, and BeFree for genes, genes, drugs, and diseases, respectively. From left to right, the *p*-values comparing MeTeOR to the literature-derived networks were 0.0076 (t=3.707, df=7), 0.0001 (t=7.822, df=7), 0.0432 (t=2.125, df=26),0.094 (t=2.145, df=5), <0.0001 (t=5.172, df=48). Excluding EVEX, MeTeOR’s difference from the literature-derived networks is still significant *p*-value<0.0005 (t=3.79, df=40). B) MeTeOR contained more known associations than the comparable algorithms in two of the three cases, and possessed more overall (17% vs 12%). C) In order to test the ability of the network to reliably predict biomedical associations, we performed a time-stamped, or retrospective, study.

### MeTeOR outperforms literature-derived databases in number and reliability of associations

To assess the coverage and quality of MeTeOR, we compared it first with specialized,gold-standard databases. MeTeOR tallies about twenty percent more gene-gene associations than BIOGRID low-throughput associations (177,000 vs 147,000; **Supplemental Fig. 3**). More impressively, MeTeOR contains 16.4 and 15.9 fold more gene-disease and gene-drug associations than CTD and DGIdb respectively. Yet despite these gains in associations, MeTeOR overlapped each of these control databases to the same extent that they overlapped each other (**Supplemental Fig. 3**).

**Figure 3.**
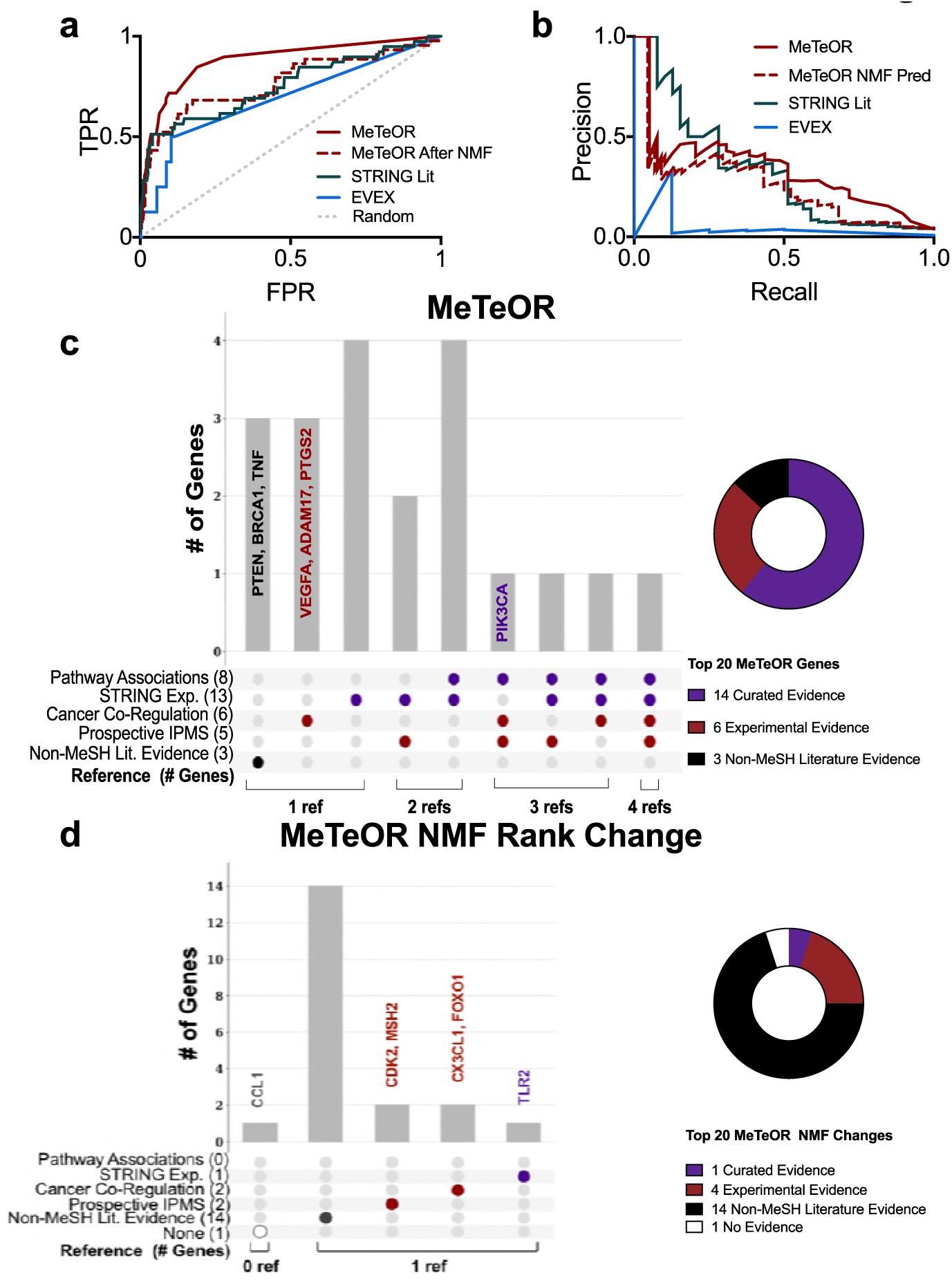
MeTeOR-identified associations with EGFR and NMF predictions. **A)** EGFR MeTeOR, STRING-Literature (lit.), and EVEX literature associations are compared against pathway-level interactions, with AUCs of 0.88, 0.75, and 0.69, respectively. **B)** In the precision recall curve, MeTeOR’s initial false positive rate is lower than that for EVEX, but higher than that for STRING-Lit. **C)** The overlap of the top 20 MeTeOR Genes with curated (MSIGDB and STRING Experimental) and experimental (Cancer Co-Expression and Prospective ImmunoPrecipitation Mass Spectrometry) evidence. Genes that did not fall into these categories were verified in the literature manually or determined to have no evidence (Supplemental Data File 2). Genes possessing experimental evidence and/or one or two references of support, which are of particular interest, are written on the chart. Genes classified with Curated Evidence have at least curated, with the possibility of Experimental or Non-MeSH Literature Evidence, with Experimental Evidence having at least Experimental. **D)** The top 20 ranked genes by their difference from MeTeOR’s rankings to their rankings after NMF were also compared against the same references. All but one of the genes (CCL1) possessed some evidence.

MeTeOR also proved as reliable as these databases, preferentially recovering the high-quality reference annotations over novel information. In other words, when MeTeOR associations were ordered by confidence (the number of supporting articles), the area under the Receiver Operating Characteristic (ROC) curves (AUC) averaged 0.71 for all references (**Figure 2A**). The average precision at 10% recall was 0.85 and at 50% recall was 0.73. (**Supplemental Fig. 4**).

**Figure 4.**
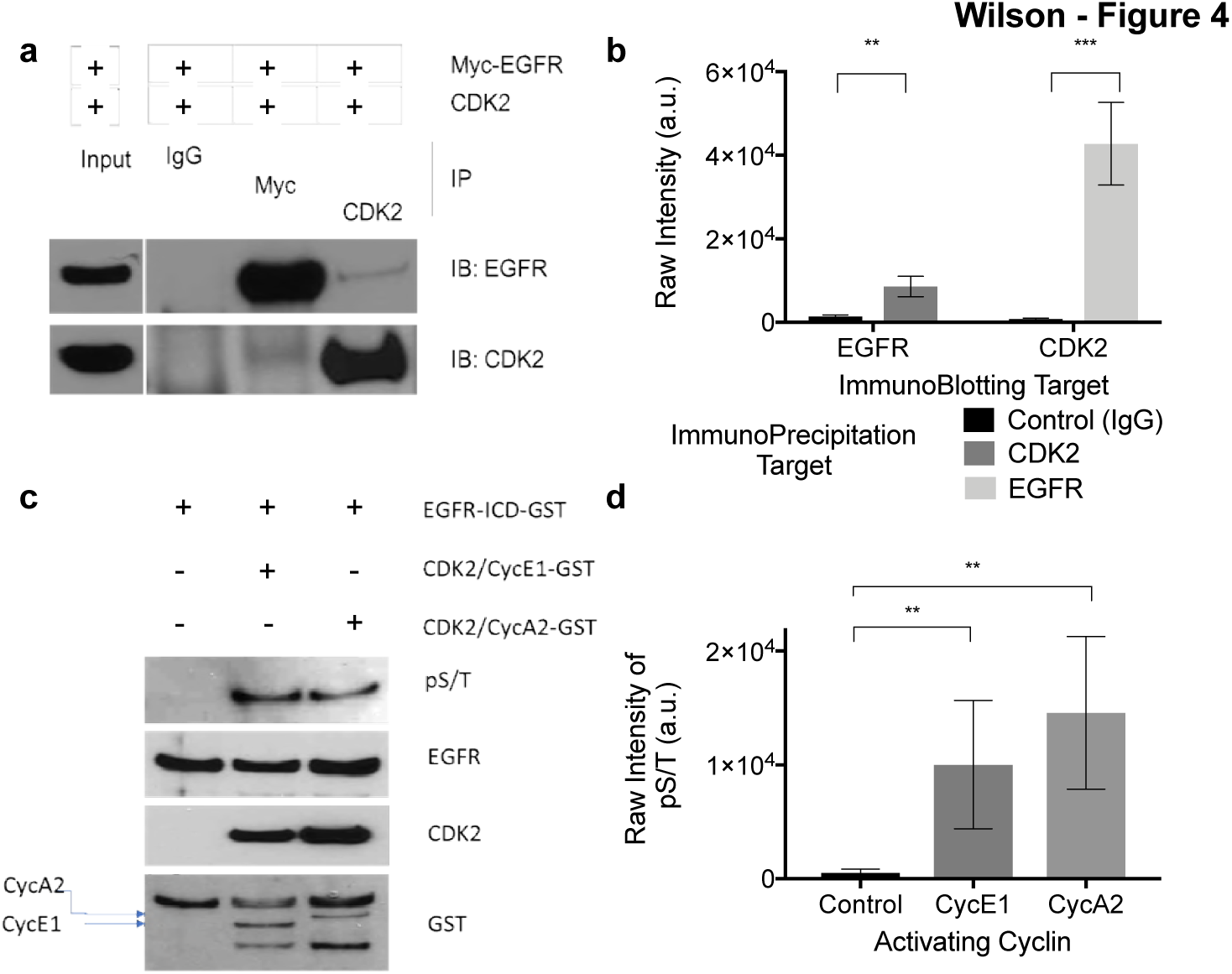
CDK2 phosphorylates EGFR, as predicted by MeTeOR. A, B) The western blot of the *in vivo* reciprocal pull-down of EGFR and CDK2 provided evidence of physical interaction between EGFR and CDK2. HEK293 cells were transfected with myc-tagged WT-EGFR and WT-CDK2 vectors, and overexpressed EGFR and CDK2 were immunoprecipitated from lysates using anti-myc or anti-CKD2 antibody and quantified over three to five replicates. Mouse IgG antibody was used as a control. C, D). An *in vitro* kinase assay showed Serine/Threonine phosphorylation on EGFR by CDK2 with statistically significant levels being generated with either Cyclin A or Cyclin E activating CDK2. Purified recombinant EGFR-GST was incubated with recombinant cyclin A2 and cyclin E1 activated CDK2 kinase; quantification on three replicates for CDK2-Cyclin E and CDK2-Cyclin A was performed with ImageJ (* *p*-value<0.05,** *p*-value<0.01, *** *p*-value<0.001; *in vivo*: t=6.834, df=4 for EGFR and t=3.407, df=8 for CDK2; *in vitro*: t=4.961, df=6 for CycA2 and t=3.984, df=6 for CycE1).

We next compared MeTeOR to the literature-mining methods STRING-Literature [35], EVEX [22], BeFree [36], and STITCH-Literature [37]. These methods extract only one type of association from the literature—gene-gene, gene-disease, or gene-drug, respectively—and MeTeOR outperformed each of them across all references, except the BeFree method on the CTD reference. MeTeOR also outperformed all methods combined, both with and without the poor-performing EVEX (**Figure 2A**). It is worth noting that MeTeOR contained several-fold more novel associations than these other text-mining tools (**Figure 2B**), even though it has roughly the same order of magnitude of overlap with the references (**Supplemental Fig. 3**). These data show that MeTeOR mines more gene-gene, gene-disease, and gene-chemical associations than are found in our reference databases, while simultaneously recovering high-quality references better than the state-of-the-art text-mining tools.

The product of the latent matrices W and H from pre-2014 data resulted in a new network with predictions. Predictions were validated if they were borne out in the literature between 2014 and 2018. D) The area under the ROC was calculated for MeTeOR gene-gene, gene-disease, and gene-drug associations based on Nonnegative Matrix Factorization (NMF) predictions being present in the 2018 network. These were compared against predictions from two naïve predictors, Common Neighbors (CN) and Adamic/Adar (AA). E) Positive predictive values (precision) were calculated at 10% and 50% recall. (* *p*<0.05, ** *p*<0.01, *** *p*<0.001).

### Testing MeTeOR predictions with retrospective analyses

We next tested MeTeOR’s ability to predict novel associations among genes, diseases, and drugs. Kastrin et al. recently tested both supervised and unsupervised link prediction methods on a MeSH co-occurrence network of 27,000 entities and found they could generate reliable hypotheses [30]. We hoped to build upon this attempt by using a more advanced link prediction method, Non-negative Matrix Factorization (NMF), with our greater number of entities (totaling 101,000). Often used in biology [38, 39], NMF is a semi-supervised machine learning algorithm that determines missing associations in a graph by decomposing it into a product of matrices [40]. Therefore, we tested the predictive power of the top two unsupervised algorithms from Kastrin et al. [30], Adamic/Adar (AA) and Common Neighbors (CN), and NMF in a retrospective study.

Here, we used cross-validation to estimate the number of features for each part of the NMF decomposed matrix (**Figure 2C, Supplemental Table 1**). When we applied NMF, we used a representation of MeTeOR derived solely from publications up to and including the year 2013 to test whether MeTeOR’s predicted associations would be confirmed by appearing in literature published between 2014 and 2018. The median AUCs of gene-gene, gene-disease, and gene-drug associations were 0.65, 0.69 and 0.67, respectively (**Figure 2D**, *left*), while the median precisions at 10% recall (the top 10% of the highest-confidence associations) were 0.75, 0.79, and 0.81, respectively and 0.65, 0.68, 0.67 at 50% recall (**Figure 2D***, middle and right*). Moreover, using AA and CN results in random predictive power, or AUCs at 0.5 (AA: 0.53, 0.50, 0.49; CN:0.51, 0.46, 0.50; for gene-gene, gene-disease, and gene-drug median AUCs, respectively). It is important to note that the AA and CN predictions are distinct from previous attempts [30] in that the network excludes many general MeSH terms, includes SCRs, and is split into separate association modes. Due to NMF’s reliable and higher performance, we chose it for subsequent analyses. These data show that the hypothetical associations among genes, drugs, and diseases produced by MeTeOR are likely to be confirmed in subsequent literature, especially those with the best confidence.

We investigated some of the top time-stamped associations in more detail in order to confirm the biological relevance of these predictions. To date, the literature has provided supporting evidence for 19, 17, and 18 out of the top 20 hypotheses from gene-gene, gene-disease, and gene-drug associations, respectively (**Supplemental Data File 1**). For example, a top predicted gene-gene association, based solely on the literature published up to and including 2013, was between the human MeSH terms for *MSX1* and *CXCR4*. In 2017, a paper was published showing that both *MSX1* and *CXCR4* independently regulate the motility and development of a population of highly migratory cells, known as primordial germ cells which give rise to eggs and sperm migration [41], and confirming MeTeOR’s hypothesis that these genes are linked in a biologically meaningful manner. To demonstrate a more complex, specific and novel prediction, MeTeOR predicted an association between *PTEN* and glaucoma based on pre-2013 literature. In the beginning of 2018, a paper was published demonstrating that microRNA MiR-93-5p, which targets PTEN, regulates NMDA-induced autophagy in glaucoma. Several other papers published after 2014 [42, 43] also suggested some role for PTEN in glaucoma. MeTeOR also predicted an association between GLI1 and multiple myeloma, and in 2017, Alu-dependent RNA editing of GLI1 was shown to promote malignant regeneration in multiple myeloma [44].

There were also some more complex indirect three-way associations (ex. gene-disease-gene). For example, the top gene-gene prediction is between CD27 and CXCR4. This prediction makes sense in the context of the human immunodeficiency virus (HIV), where HIV-1 variants use CXCR4 to infect T cells, and through this process, HIV depletes both naïve and CD27^+^ memory T cells [45]. This demonstrates the predictive power of the network by highlighting a complex gene-disease-gene relationship (CXCR4 – HIV – CD27). Another example is between *WT1* and *HLA-B*. The WT1 protein has been chosen as an immunologic target by a National Cancer Institute initiative [46], and this year, a phase 2 clinical trial showed a WT1 vaccine that is effective in Acute Myeloid Leukemia with predicted binding on HLA-B*15:01, HLA-B*39:01, HLA-B*07:02, and HLA-B*08, HLA-B27:05 in addition to HLA-A*02 [47]. These MeTeOR predictions suggests that further investigation is warranted and highlights the ability of the network to suggest complex gene–disease–gene relationships.

Though these hypotheses are only a small sample of all MeTeOR-identified links, they illustrate the power and range of MeTeOR’s NMF predictions.

### MeTeOR identifies known and novel *EGFR* associations

To illustrate how MeTeOR might be used, we focused on Epidermal Growth Factor Receptor (EGFR) as a test case. EGFR is a well-studied protein involved in various aspects of carcinogenesis [48], and we hypothesized that MeTeOR would be able to extract known and novel associations from the wealth of extant literature.

We first needed to understand EGFR’s known and verifiable associations. MeTeOR found 1064 genes connected to EGFR via MeSH terms in at least one article, 467 genes in at least two articles, and 97 genes in at least ten articles. Assuming that associations made by more articles would be more robust, we compared the MeTeOR-ranked list of 1064 gene-EGFR associations against the MSIGDB pathway standard used in Figure 2.

MeTeOR recovered pathway information better than the text-mining algorithm EVEX (overall AUC_MeTeOR_ of 0.88 *vs* AUC_EVEX_ of 0.69; **Figure 3A**). MeTeOR’s initial recall was also superior, as indicated by the Precision-Recall curve (**Figure 3B**). Finally, MeTeOR was overall more accurate than STRING Literature (AUC_STRING_ of 0.75), although in the initial recall, STRING did better, likely because it weighs confidence based on KEGG pathway information [49] (**Figure 3B**).

We then sought to evaluate MeTeOR’s likelihood of generating false positives. Reliance on MeSH terms could, for example, create a spurious link between EGFR and another gene if the publication is a review article that mentions another gene without actually proposing a relationship with EGFR. We noticed that 12 of the top 20 genes MeTeOR associated with EGFR did not appear in MSIGDB pathway standard (**Figure 3C, Supplemental Fig. 5**). We therefore compared these top 20 genes against experimental associations derived from public sources (aggregated in STRING-Experimental). The STRING-Experimental dataset (STRING-EXP), which showed that 13 out of the top 20 genes physically interact with EGFR (**Figure 3C**), revealed that six of the twelve genes missed by MSIGDB are actually valid (**Supplemental Fig. 5**). This brought the number of genes with curated evidence from MSIGDB pathways or STRING from eight up to 14 (**Figure 3D**). For the remaining six genes, we pursued two analyses based on experimental evidence, one involving pan-cancer RNA-seq data (from 8768 TCGA patients [50], see Online Methods) and the other a prospective, unbiased high-throughput Mass Spectrometry experiment.

We calculated the co-expression of all genes in 20 TCGA cancer types and thresholded them by the correlation co-efficient. The mRNA levels of three of the six putative “false positive” genes correlated with EGFR mRNA levels (|r| >0.25, Online Methods). For example, PTGS2 was not associated by pathways but was co-expressed with a q-value << 0.01, r = 0.29. This appears to be a biologically relevant relationship insofar as both PTGS2 and EGFR are prognostic biomarkers for several of the same cancers [51, 52], and PTGS2 expression levels can predict the efficacy of treatments that act on EGFR [53]. EGFR associations with the other two genes (*VEGFA* and *ADAM17*) appear equally valid (**Supplemental Data File 2**).

For the high-throughput Immuno-Precipitation Mass Spectrometry (IPMS), we pulled down EGFR at several time-points after stimulation with Epidermal Growth Factor (EGF) in order to obtain a snapshot of proteins binding with EGFR in a functional context (**Supplemental Fig. 6, Supplemental Data File 3**). IPMS showed that five of the 20 genes were associated with EGFR, though all were also associated with MSIGDB pathways or STRING. One of these five was *PIK3CA*, which possesses links through pathway knowledge, cancer co-regulation and the IPMS; it is frequently co-mutated with EGFR [54] and known to interact with other PI3K subunits (PIK3CB [55] and PIK3R1 [56]) [57].

In the end, just three genes (*PTEN*, *BRCA1*, and *TNF*) remained putative false positives (**Figure 3C**). All three, however, have some degree of literature support, denoted as non-MeSH literature evidence because it is manually curated and not originating from MeSH terms (**Supplemental Data File 2**). For example, *PTEN* is often lost in cancers with *EGFR* gains [58] and the EGFR/PI3K/PTEN/Akt/mTORC1/GSK-3 pathway causes malignant transformation, drug resistance, metastasis, and prevention of apoptosis [59]. Thus, even the apparent false positives in the top 20 associations seem to warrant investigation.

### MeTeOR’s automated hypothesis generation predicts new EGFR associations

Although the success of MeTeOR’s retrospective associations is reassuring, the real test of MeTeOR’s utility to the scientific community is whether it can reveal unexpected and valuable biological hypotheses that merit experimental validation. We therefore used EGFR as a test case again, but instead of using MeTeOR’s raw associations, this time we evaluated its Non-Negative Matrix Factorization (NMF) predictions. These were ranked by their difference from MeTeOR’s rankings, such that: *NMF Rank Change* = *MeTeOR Rank* − MeTeOR *NMF Rank* where *MeTeOR* Weight>2 limits arbitrarily large ranks from genes that initially had little to no evidence (**Supplemental Data File 4**).

Controlled against MSIGDB pathway associations, all 20 predictions were putative “false positives” and only one possessed STRING-Experimental evidence (TLR2) (**Figure 3D**). This demonstrates the effectiveness of the NMF Rank Change at highlighting novel predictions. Yet, of the 19 unproven associations, two were co-expressed in cancer (CX3CL1 and FOXO1) and two were supported by our IPMS evidence (CDK2 and MSH2) (**Supplemental Fig. 5**). Of the remaining fifteen genes, all except CCL1 had non-MeSH literature support (**Figure 3D; Supplemental Data File 2**), underscoring the quality of NMF Rank Change predictions.

To narrow down candidates for experimental validation, we focused on CDK2 and MSH2, the proteins for which we had IPMS evidence (**Figure 3D**). Cyclin-dependent kinase 2 (CDK2) seemed the most biologically promising: like EGFR, CDK2 is directly involved in the cell cycle and cell growth, and it has a similar kinase domain to CDK1, which phosphorylates EGFR *in vitro* [60]. Furthermore, in apoptosis and senescence, CDK2 translocates to the cytoplasm with Cyclin A [61] or Cyclin E [62], and under these conditions, an activated CDK2 might bind to and phosphorylate EGFR.

To determine whether CDK2 and EGFR directly interact in a biologically relevant manner, we transfected human embryonic kidney cells with expression vectors for both proteins. Co-immunoprecipation demonstrated that CDK2 and EGFR formed stable protein-protein interactions (**Figure 4A, B**). Next, we incubated purified EGFR protein by itself or with CDK2, along with either its interaction partner Cyclin A2 or Cyclin E1. We found that, *in vitro*, both Cyclin A and Cyclin E activate CDK2 to phosphorylate EGFR’s intracellular regulatory portion (**Figure 4C, D**) but not to phosphorylate the extracellular portion (**Supplemental Fig. 7**). *In silico* prediction with GPS [63] identified several residues (752, 847, 991, 1026, 1032, and 1153) as possible sites of intracellular EGFR phosphorylation by CDK2 (**Supplemental Fig. 8**). It is worth noting that Residue 1026 was previously shown to be phosphorylated by CDK1 [60].

This interaction is rather surprising because CDK2 has never been shown to interact with EGFR. Yet our data indicate that CDK2 directly phosphorylates EGFR, and they bind to one another *in vivo.* MeTeOR’s automated hypothesis-generation thus produced many validated biological hypotheses, and in the case of CDK2 has revealed an unexpected and valuable biological insight.

## DISCUSSION

Our ability to find interesting relationships among bodies of knowledge separated by time and disciplinary boundaries is struggling with the ever-increasing size of the scientific literature [1]. Current tools, such as PubMed and Google Scholar, make it possible to search extant publications (at least to the extent that the content is available online), but they can reflect and propagate biases [64]; they cannot evaluate the relative confidence of observations; and they do not attempt to integrate information into novel hypotheses. Whereas many literature-mining methods seek to capture semantic and syntactic detail from each paper, we took the opposite approach, hypothesizing that millions of human-curated keywords could create useful network structures and that the sheer quantity of data points would wash out erroneous results while allowing verifiable information to emerge from separate but corroborating studies. Following the Bag-of-Words representation of knowledge in terms of common, contextual word associations [65], we focus on the most important facts from each paper embodied by (key) words chosen from Medical Subject Heading (MeSH) terms. These MeSH terms are readily available and regularly updated. By representing each article as a clique of MeSH terms, we create networks that can reveal unsuspected connections across the literature. This effectively converts unstructured into structured knowledge that, in turn, is amenable to machine learning techniques to generate new hypotheses.

In practice, the MeSH Term Objective Reasoning (MeTeOR) network pooled knowledge from over 22 million PubMed articles to create a map of relationships among genes, drugs, and diseases. MeTeOR recovered knowledge from reference databases and revealed many previously uncharacterized biomedical associations; its performance was on par with or better than domain-specific and state-of-the-art Natural Language Processing (NLP) models for knowledge extraction. Moreover, hypothesis generation through non-negative matrix factorization predicted new associations prior to their publication. This predictive efficacy was further demonstrated by MeTeOR’s ability to discern known and novel EGFR interactions more reliably than NLP algorithms. In particular, MeTeOR predicted an association between CDK2 and EGFR, and we confirmed and simultaneously suggested the association is a direct physical interaction with high-throughput IPMS screening. This interaction has implications for biological processes such as cell cycle, cell growth, and apoptosis as well as disease processes such as tumorigenesis. Both CDK2 [66] and EGFR [67] are targets of cancer therapies, but previous hints of a relationship between the two proteins had been attributed to similarities in structural activation [68] or distant regulatory effects [69]. Our experimental data verified this interaction, which had been latent in the literature but gone unnoticed. Together, these results demonstrate that the breadth and redundancy of keyword coverage in the literature compensate for the superficiality of the information taken from any one article and can accurately represent knowledge across a large corpus of literature, creating hypotheses that warrant experimental investigation.

In the future, MeTeOR can be improved in a number of ways. It could be combined with orthogonal databases [49] or ontological hierarchies [70] so as to improve the network accuracy and coverage. Additional relevant keywords, such as the context of an association (e.g., regulation, phosphorylation) and MeSH terms for biological processes, therapies, and clinical variables, could deepen MeTeOR analyses. Labels that convey dates, number of citations, journal, and other contextual details might provide useful qualifiers for the confidence of associations. Alternatively, defining the semantic meaning of the relationship may be done through integration of the SemRep system [71]. Keyword indexing exists in fields outside biomedicine [72] and could be turned, likewise, into knowledge networks that summarize and support machine learning over entirely different domains of knowledge. For now, MeTeOR is a public, reliable source of gene, drug, and disease associations that directly link to PubMed references, improving accessibility and indexing of the literature, while enabling its use for hypothesis generation across biology.

## MATERIALS AND METHODS

### Indexing Information to Represent Biomedical Knowledge

Co-occurrence strengthens the confidence in associations as the number of articles sampled increases [73]. Supplementary Concepts Records (SCRs) are similar to MeSH terms and cover a wide variety of concepts including genes, drugs, and diseases. They were used in addition to MeSH terms to supplement the existing data. All data was obtained using the NCBI eutils tool and a list of all PubMed IDs associated with a search for Eukaryotes, Bacteria, Viruses, and Archea (∼22 million articles). All proteins were mapped to Entrez ids using supplementary concepts annotations of RefSeq numbers in the notes section where possible and by symbol or synonym if no RefSeq number was present. All drugs and diseases were mapped using the MeSH hierarchy as done in previous works [21, 74], with PubChem CIDs used for drugs and MeSH ids for diseases. In order to obtain the co-occurrence of these terms, we calculated the dot-product of the term-article membership matrix. Terms that mapped to the same Entrez ids were summed by edge weights.

### Data Visualization

The MeTeOR network was filtered to only use edges that had a confidence over 200, and while nodes were made invisible. The weights of each edge represented as the penwidth for each edge. The format for the network was assembled in NetworkX (https://networkx.github.io/) in python as DOT file, and then the network was visualized using the sfdp tool of GraphViz (http://www.graphviz.org/).

### Ground Truth Comparisons

The network was compared against highly accessed and cited databases in order to determine if the network contains valid associations between terms. These comparisons measure the recovery of a reference database based on the ranking of the others (MeTeOR or a literature-derived source), and the data output is the recovery rate of true positives (TPR) and false positives (FPR). A true positive was defined as an association present in MeTeOR that also was present in the ground truth.

### Robust Comparisons

Receiver Operating Characteristic (ROC) plots can lead to inaccurate representations of the data when there are unbalanced numbers of true and false negatives. In particular, if there is a space of 100,000 by 100,000 possible associations between drugs and genes, most of the possible interactions will be True Negatives, making the False Positive Rate increase extremely slowly according to the formula:

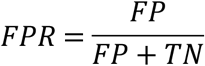

This leads to inflated AUCs. To solve this problem, the number of positives and negatives was determined, and an approximately equal number of positives and negative were chosen randomly together up to a hundred times. This was designed to randomly sample for complete coverage of all positives. Occasionally, the number of positives per iteration was below 100, and in order to make each iteration more reliable, the number of iterations was decreased. This allowed the determination of a range of accuracy scores (ROC, PR, etc.) for each comparison. The final comparison between MeTeOR and a literature-derived source was calculated with a paired t-test on the group of average AUCs or PRs from the bootstraps. Any reference which had fewer than 3 overlaps with either MeTeOR or a literature-derived source was discarded. Additionally, references were broken down by type if provided (example: BIOGRID High and Low Throughput).

### Box Plots and Statistics

Boxes define the 25^th^ -75^th^ percentiles, with the whiskers extending from min to max, and the line in the middle defining the median. All statistical tests are two-sided. For comparisons against the ground truths in Figure 2A, all values are means of the bootstrap values, and these means were compared with a paired t-test, when all values were pooled together, they passed a D’Agostino & Pearson normality test with a K2=1.615, p=0.4459 for the literature-derived source and K2=0.6366, *p*=0.7274 for MeTeOR.

### Data Normalization

The MeTeOR network was smoothed using Laplacian normalization, as defined by:

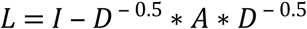

where L is the normalized Laplacian, D is the degree matrix, and A is the adjacency matrix of the network. This was done for each mode (gene-gene, gene-disease, gene-drug, etc.). For large-scale ranking, the absolute value of the non-diagonal elements was used. In individual rankings, such as to EGFR, the non-normalized data was used to provide easy interpretation.

### Collection of Ground Truths

In order to determine if MeTeOR contained valid gene, disease, and drug information, ground truths were collected from the literature. MSIGDB refers to the canonical pathways from MSigDB [75] and was used to determine gene-gene pathway-level associations, while the components of BIOGRID [19] represented physical gene-gene associations. A gene-gene association was made for MSIGDB if two genes were present in a pathway together, and each association was given a confidence 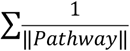 and then all confidence scores were normalized to Z-scores. The top 0.1% of associations (N=32,000) were used as a ground truth to prevent promiscuous associations.

There were several databases for gene-disease associations including the Comparative Toxicogenomic Database (CTD) [20] and DisGeNET [76], and these databases were broken down into their component pieces and mapped to Entrez IDs for genes and MeSH terms for diseases. For gene-drug interactions, the primary sources of data were DGIdb [77] and Drugbank, downloaded through BIOGRID [19]. Pubchem CIDs [32] were used to map MeSH chemicals [32] and Drugbank’s mapping facilitated Drugbank IDs to CIDs. All STRING networks were mapped to Entrez IDs though STRING’s provided mappings from STRING 9 and STRING 10. All references were retrieved in March 2018. Mappings created in this project can be found within the data repositories provided with this paper.

### Collection of Text-Mining Algorithms

STRING-Literature (version 10.5), EVEX, STITCH-Literature (version 5), and DisGeNET’s BeFree (version 5) were chosen as representative Natural Language Processing (NLP) efforts to mine gene-gene, gene-drug, and gene-disease relationships from the literature. All these efforts are publicly available and have been through multiple revisions as they undergo continued development.

### Naïve Unsupervised Prediction Methods

Two naïve methods were used to compare against a more advanced algorithm, Non-negative Matrix Factorization (NMF). These algorithms were the Common Neighbors algorithm and the Adamic/Adar algorithms, calculated to include edge weight confidence. These were selected because of their top performance in Kastrin et al.[30].

Though it is worth noting that in this publication, we include SCRs and limit the analysis to specific edge types (e.g. gene-gene), which is not true in Kastrin et al.[30].

### Non-negative Matrix Factorization (NMF)

The principle behind NMF is to create two low-dimensional matrices that, when multiplied together, approximate an original matrix [40]. These matrices are called basis vectors, where the degree to which they can recapitulate the original matrix is determined by their size. The greater the size, the more features the basis vectors can capture. The basis vectors are determined through several optimization algorithms that act upon randomly initialized W and H matrices. In this work, we employed both the alternating least squares algorithm:

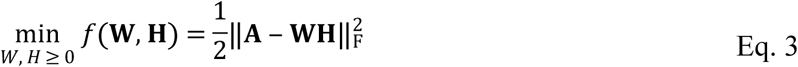

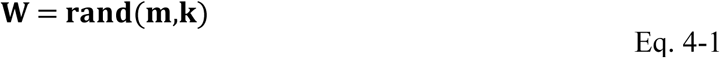

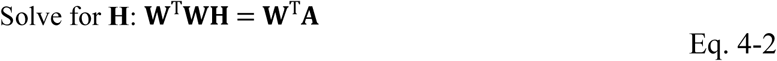

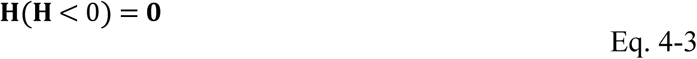

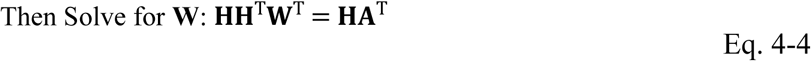

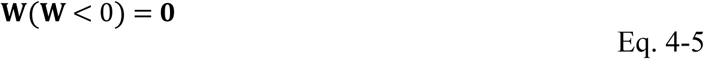

and the multiplicative algorithm:

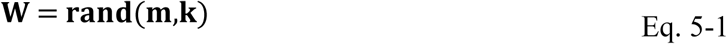

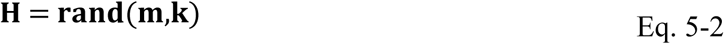

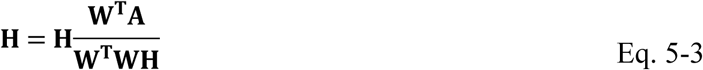

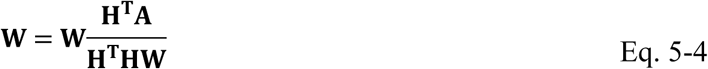

NMF was executed computationally with MATLAB’s Statistics Toolbox, with three repetitions of 5 iterations of the multiplicative algorithm in order to find the optimal basis initialization, then 100 iterations of the alternating least squares were performed. For bulk analysis, this was done one time. For specific association predictions, like associations to EGFR, this NMF process was completed five times, and then the Mean Reciprocal Rank was computed for each association across the NMF runs. This ensured that a stable answer was obtained despite the non-convex nature of NMF. The number of features (k) was selected using ten-fold cross validation of each mode of MeTeOR. The Matthew’s Correlation Coefficient (MCC) was calculated and rounded to two digits of significance in order to select the lowest k with the highest MCC: 300 for gene-gene, 100 for gene-disease, and 50 for gene-drug.

### Retrospective

Retrospective experiments were undertaken in order to determine if the information in MeTeOR through 2013 was sufficient to make accurate predictions that had yet to be discovered. The first retrospective experiment was a validation of the technique and quality of data, in that the MeTeOR network through 2013 was used to predict itself in 2018. After predictions were made on MeTeOR, all shared associations in the ground truth up to 2013 were removed, and the remaining predictions were assessed against the ground truth in the future.

### Tissue Culture and Crosslinking for IPMS

Hela cells were grown in DMEM (Sigma) with 10% FBS (Invitrogen) in 5% CO2 at 37°C. 10^8^ cells were crosslinked with formaldehyde by directly adding it to the culture medium to a final concentration of 0.5% for 8 min at 37°C. The cross-linking reaction was quenched by adding Glycine (Sigma) to a final concentration of 0.2M. Membrane proteins were extracted by re-suspending the pellet in LB1 buffer (50mMHEPES-KOH [pH 7.5], 140mMNaCl, 1mMEDTA, 10% glycerol, 0.5% NP-40, 1% Triton X-100) for 30 min at 4°C. After centrifugation the supernatant containing crosslinked membrane and cytosolic proteins was used for immunoprecipitation. Immunoprecipitation and sample prep for mass spectrometry was performed as previously described [78].

### Mass Spectrometry

Binding partners of EGFR were pulled down at different time points (2, 10, 30, 120 seconds) after EGF stimulation and identified through ImmunoPrecipitation Mass Spectrometry (IPMS) in HeLa cells. Each IPMS experiment was conducted in triplicate, with one IPMS experiment conducted on non-stimulated cells to serve as a baseline. Peptides were reconstituted in 0.5% methanol, 0.1% formic acid and fractionated using a C18 (2 µm, Reprosil-Pur Basic, 6 cm x 150 µm) column with an EASY-nLC-1000 HPLC (Thermo Scientific) online with a Q-Exactive mass spectrometer (Thermo Scientific). A 75-minute gradient of 2-26%acetonitrile, 0.1% formic acid at 800nl/min was used per fraction. A window of 300-1400 m/z at 120k resolution, 5 × 10^5^ AGC, and 50ms injection time, was used for precursor selection. The top 50 most intense ions were selected for HCD fragmentation with a 5 m/z isolation window, 18 sec exclusion time. RAW files were acquired with Xcalibur (Thermo) and processed with Proteome Discoverer 1.4 and MASCOT 2.4. Peptides were matched using a 20 ppm precursor tolerance window and 0.5 Da fragment threshold. Up to two missed cleavages were allowed. The data was filtered with a 1% false discovery rate by Percolator and abundances were calculated by the iBAQ algorithm. RAW files were then converted to mzXML and peptide abundances were distributed to gene products through Grouper software. Unique to gene PSMs must be >=1.

### Analysis of Mass Spectrometry

All EGFR-associated proteins had their iBAQ levels normalized across time points and averaged across three biological replicates. All missing values were filled in with the minimum overall value. The amount at a given time point was calculated as a gradient relative to the previous time point. The gradient allowed the monitoring of protein changes over time, and clustering of the gradients through k-means revealed distinct patterns (Supplemental Fig. 6). Most patterns were self-consistent and showed a change at the initial time points, with little change thereafter, but the second group appeared to show random changes for proteins over all time points and may be promiscuously associated with EGFR (Supplemental Fig. 6). All proteins that changed more than 5% over the course of the experiment were considered true positives and associated with EGFR.

### *In vitro* Kinase Assay

Two hundred fifty ng of purified recombinant EGFR-GST (Aa 668-1210, Sino Biological Inc, Beijing, P.R. China) was incubated with 100 ng of recombinant cyclin A2 or Cyclin E1 activated CDK2-GST kinase (ProQinase, Freiburg, Germany) in 20 µl of kinase buffer (10 mM HEPES, pH 7.5, 50 mM glycerophosphate, 50 mM NaCl, 10 mM MgCl2, 10 mM MnCl2, 1mM DTT and 10 µM ATP) for 30 min at 30°C. The reaction was terminated by addition of SDS treatment buffer, applied to 4-12 % SDS-PAGE, and immunoblotted with anti-phopho-S/T (BD Bioscience, San Jose, CA, USA), anti-EGFR, anti-CDK2, or anti-GST antibodies (Santa Cruz Biotechnology, Dallas, TX, USA).

### *In vivo* Reciprocal Pull-Down

HEK293 cells were grown in 6 cm dishes and transfected with 2 µg of WT-EGFR and WT-CDK2 expression construct using lipofectamine 2000 (Life Technologies, Carlsbad, CA. USA). After 24 h incubation at 37°C, cells were lysed with Buffer (10 mM HEPES, pH 7.5, 10 mM KCl, 0.1 mM EDTA, 1 mM DTT, 0.25% NP-40) containing protease inhibitor cocktail (Roche). Lysates were centrifuged at 6,000 rpm for 4 min and the supernatants were transferred to a new tube and protein concentration measured using Bradford assay (Bio-Rad Laboratories, CA). One hundred µg of cell lysate was incubated with 2.5 ug of anti-myc antibody (BioLegned, San Diego, CA, USA) or anti-CDK2 antibody (Santa Cruz Biotechnology, Dallas, TX, USA) overnight at 4°C. After further incubation with 20 µl of protein A agarose (50% (v:v) in lysis buffer (Santa Cruz Biotechnology, Dallas, TX USA), the incubation mixture was washed three times with 1 ml lysis buffer, and twice with RIPA buffer (Boston BioProducts, MA, USA) containing protease inhibitor cocktail V (Calbiochem, CA, USA). The precipitates were re-suspended in 20 µl of 2 × SDS sample buffer and heated at 100 °C for 5 min and were applied to 4-12% SDS-PAGE followed by immunoblotting using anti EGFR, anti-CDK2, or anti-GST antibodies (Santa Cruz Biotechnology, Dallas, TX, USA). Mouse IgG antibody (Santa Cruz Biotechnology) was used as a control.

### Co-Regulation of Genes in Cancer

The RNASeqV2 Level 3 files of 20 TCGA cancer types (BLCA, BRCA, CESC, COAD, GBM HNSC, KIRC, KIRP, LAML, LGG LIHC, LUAD, LUSC,OV PRAD, READ, SKCM, STAD, THCA, UCEC) were downloaded from TCGA data portal (https://tcga-data.nci.nih.gov/tcga/) on August 19, 2015. RSEM (RNA-Seq by Expectation Maximization [79]) normalized count values of 8,768 tumor samples were used to compute Spearman’s rank correlation coefficient of EGFR and all other 20,426 genes. Genes with absolute values of correlation coefficient more than 0.25 were considered to be significantly co-regulated with EGFR.

### EGFR NMF Predictions

Because the Non-negative Matrix Factorization (NMF) predictions are based on MeTeOR associations, the NMF MeTeOR rank was subtracted from the MeTeOR rank, to obtain a MeTeOR Difference.

### Data and Code Availability

All data and code from the MeTeOR network is available online at http://meteor.lichtargelab.org/ or http://osf.io/as865.

### Computation

MeTeOR was assembled in python 3 and tested using MATLAB code for comparisons on an Ubuntu computer with 64 GB RAM and 4^th^ Gen. Intel Core i7 3.7 GHz processor.

## ACKNOWLEDGEMENTS

The authors thank Vicky Brandt, Christie Buchovecky, Teng-Kui Hsu, Rhonald Lua, and Panos Katsonis for their discussions and general feedback on the work.

## AUTHOR CONTRIBUTIONS

SJW conceived of the project, designed the experiments, wrote the code for the experiments, and wrote the manuscript. ADW formed the initial interest around MeSH terms, helped guide the experiments and edited the manuscript. MH helped interpret and process the IPMS experiments, which YC conducted, with YI and JQ overseeing. BKC conducted the *in vitro* and *in vivo* experiments overseen by LD. DK, CHL, AK, and BW helped with experimental design and manuscript preparation. CHL prepared the TCGA RNA-Seq data. SYK helped design and implement the website. OL oversaw all experiments and manuscript preparation.

